# *Cryptosporidium* oocysts and *Giardia* cysts at two South African wastewater treatment plants

**DOI:** 10.1101/2024.07.22.604670

**Authors:** Thandubuhle Gonose, Hein Du Preez, Stephanus N Venter, Monique Grundlingh

**Affiliations:** Rand Water Analytical Services, PO Box 3526, Vereeniging, South Africa; University of Pretoria, Department of Microbiology and Plant Pathology, Pretoria

**Keywords:** Prevalence, wastewater treatment, *Giardia*, *Cryptosporidium*, wastewater treatment plants

## Abstract

*Cryptosporidium* and *Giardia* are one of the most prominent parasitic protozoans found in wastewater treated by wastewater treatment plants (WWTPs). They cause Cryptosporidiosis and Giardiasis respectively, which can be fatal in immuno-compromised individuals. Thus, their removal by WWTPs is essential. This study aimed at investigating the prevalence of *Crytposporium* and *Giardia* through treatment processes at two South African wastewater treatment plants and assessing the efficiency of the WWTPs in removing them. A total of 162 samples (54 raw influents, 36 primary treated effluents, 36 secondary treated effluents and 36 final treated effluents) were collected from two South African WWTPs over a period of one year. Results obtained from the study show that *Cryptosporidium* was detected in 98% (159/162) of the samples collected, whilst Giardia was detected in 99% (160/162) of the samples collected from both wastewater treatment plants. Both *Crytposporium* and *Giardia* were detected in 100% (n=54) of raw influent samples. The same result was obtained in primary treated effluent, where both *Cryptosporidium* and *Giardia* was detected in 100% (n=36) of the samples collected. In secondary treated effluent, 36 samples were collected and *Cryptosporidium* was detected in 94% of the samples, whereas *Giardia* was detected in 100% of the samples. In the final treated effluent samples, both *Cryptosporidium* and *Giardia* where detected 97% (n=36) of the samples collected. Detection of *Cryptosporidium* and *Giardia* in most of the wastewater treatment stages indicated these parasites are prevalent in the two WWTPs. Although the two WWTPs showed high removal of the two parasites in wastewater, their presence in the final treated effluent indicated the inefficiency of treatment processes in their removal. This highlights the potential of these WWTPs in contaminating receiving water sources.

**IMPORTANCE OF THE STUDY:** This study will specifically evaluate the removal of Cryptosporidium oocysts and Giardia cysts by two South African WWTPs and will estimate their ability of removing them. The study will also attempt to give insight into the potential and extend of treated effluent in polluting surface water resources with respect to Cryptosporidium and Giardia. Identification of the potential sources of the pollution is one of the significant parts in resolving this problem. Hence the finding of the study are very important in an endeavour to eliminate pollution of water source.

## 1. INTRODUCTION

*Cryptosporidium* and *Giardia* are protozoan parasites that infect the gastrointestinal tract of a broad range of vertebrate animals such as birds, reptiles, mammals, amphibians and fishes (Putignani and Menichella, 2010). In humans, *Cryptosporidium* and *Giardia* infect the distal and proximal regions of the small intestines and multiply within the hosts (Bukhari and Clancy, 2000). The infection result into a disease called Cryptosporidiosis (*Cryptosporidium*) and Giardiasis (*Giardia*) which is characterised by watery diarrhoea that could be accompanied by stomach cramps, nausea and fever (Health Canada, 2012). Infected humans excrete large amounts of *Cryptosporidium* and *Giardia*, in transmissive stages called oocysts and cysts respectively, into the environment via their feacal excretion (Putignani and Menichella, 2010).

Feacal excretion of large numbers of *Cryptosporidium* oocysts and *Giardi*a cysts by infected hosts makes these parasites some of the most frequently isolated parasites from wastewater (also referred to as sewage) (Fayer et al. 2000). Furthermore, *Cryptosporidium* oocysts and the *Giardia* cysts excreted by the infected hosts are very robust and can remain viable under different environmental conditions for long periods. They are also able to resist oxidizing disinfectants typically used by water suppliers to remove pathogens (Carmena, 2010). Because of these features, *Cryptosporidium* and *Giardia* are amongst the most important pathogens in the water industry.

Several studies have isolated both *Cryptosporidium* and *Giardia* from raw and treated wastewater at wastewater treatment plants in different parts of the world (Sroka et al. 2013; Castro-Hermida et al. 2010; Cheng et al. 2009; Dungeni and Momba, 2010). Isolation of the two parasites in raw wastewater could be an indication of the infection of the population by *Cryptosporidium* and *Giardia* (Robertson et al. 2000). Although high counts of *Cryptosporidium* and *Giardia* in raw wastewater is important, their detection through the treatment process and in the final treated wastewater that is discharged into water sources is of serious concern as it highlights the risk to public health. This is because the treated wastewater is discharged into surface water sources such as dams and rivers which are sometimes used by consumers for different purposes such as drinking, recreation and agriculture (Teklehaimanot et al. 2015).

Frequent detection of *Cryptosporidium* oocysts and *Giardia* cysts in the final treated effluent has shown that a number of WWTPs are not equipped to effectively remove *Cryptosporidium* oocysts and *Giardia* cysts from wastewater (Cheng et al. 2009). Various studies have confirmed this as both *Cryptosporidium* oocysts and *Giardia* cysts are regularly detected in the treated wastewater effluent that is discharge by WWTPs into water sources (Cheng et al. 2009; McCuin and Clancy, 2006; Dungeni and Momba, 2010; Robertson et al. 2006).

Factors that contribute to the inefficiency of WWTPs in removing *Cryptosporidium* oocysts and *Giardia* cysts include poor management of WWTPs, the use of ineffective treatment processes by the WWTPs and uncontrolled wastewater inputs (Nikiema et al. 2013). These factors together with financial challenges play a major role in the failure of WWTPs to produce effluent that is free of pathogens, particularly in developing countries. South Africa (SA), as a developing country, is also facing the problem of failure of WWTPs to produce effluent of high quality (Dungeni and Momba, 2010; Naidoo and Olaniran, 2014; Sroka et al. 2013). In actual fact, in South Africa WWTPs have been identified as one of the major polluters of water sources (Dungeni and Momba, 2010; CSIR, 2010).

To date, there are relatively few studies in South Africa that have looked at the prevalence and efficiency of wastewater treatment plants in removing *Cryptosporidium* oocysts and *Giardia* cysts. Thus this current study aimed at investigating the prevalence of *Cryptosporidium* oocysts and *Giardia* cysts in raw influent, through treatment stages, and in final treated effluent of two South African WWTPs over a period of 12 months. Furthermore, the removal efficiency of *Cryptosporidium* oocysts and *Giardia* cysts by the two WWTPs was evaluated. Results obtained in this study will be useful to water utilities as it can assist in the evaluation of WWTPs as polluters of the receiving water bodies and also in establishing *Cryptosporidium* and *Giardia* risk status in water sources (Sigudu et al. 2014).

## 2. MATERIALS AND METHODS

### Study sites

Two wastewater treatment plants (WWTPs) were selected as sampling sites for this study. These wastewater treatment plants were referred to as WWTP 1 and WWTP 2 and they are located in two South African Provinces namely Gauteng and the Free State. Table 1 summarises the characteristics of the two WWTPs.

**Table 1:** Characteristics of the sampling sites.

| WWTP | Wastewater source | WWTP capacity (ML/day) | Treatment processes |
| --- | --- | --- | --- |
| WWTP 1 | Domestic, Industries Abattoir | 36 | Coarse and fine steel filters, Grit remover, Settling tanks, Biological nutrient removal; Settling tanks, Chlorine dosage |
| WWTP 2 | Domestic | 2.1 | Course steel filters, Sedimentation (Settling ponds),Trickling Filters; Sedimentation; Chlorine dosage, |

*Note: WWTP1- has two raw influent receiving points. One that receives wastewater from industries and* *abattoirs and other that receives from domestic.*

### Sample collection for *Cryptosporidium* oocysts and *Giardia* cysts analysis

Grab wastewater samples were collected at different treatment stages, namely: raw influent, primary treated effluent, secondary treated effluent and the final treated effluent over a period of 12 months (February 2014 to January 2015) from WWTP1 and WWTP2. For the first six months of the study, samples were taken twice a month (second and fourth week of each month), and in the last six months sampling was done once every month. Wastewater samples were collected using 500 mL bottles for *E. coli* analysis, 1 L sampling bottles for *Cryptosporidium* and *Giardia* analysis in raw influent and 10 L sampling bottles *Cryptosporidium* and *Giardia* analysis in primary effluent, secondary effluent and final treated effluent samples.

### Sample analysis

The United States Environmental Protection Agency method 1693 was used for detecting and enumerating *Cryptosporidium* oocysts and *Giardia* cysts in the collected wastewater samples (USEPA, 2014).

#### Sample concentration

Treated effluent samples (10 L) were filtered using Pall Envirochek^TM^ High Volume (HV filter) filters at a flow rate of 2 L min^-1^. Laureth 12 elution buffer and wrist actions shakers were used to elute the filters. The eluates were transferred into 250 mL conical tubes. centrifuged using a Sorvall RC3 centrifuge at 1500 xG for 15 minutes. After centrifugation, the pellet was used to detect *Cryptosporidium* oocysts and *Giardia* cysts using the immuno-magnetic separation. Raw influent samples were directly poured into the 250 mL conical tubes and centrifuged.

#### Immunomagnetic separation

The IMS procedure was performed as described in USEPA 1693 (USEPA, 2014. Briefly, kaolin (0.75 g) was added into the LT tubes containing SL-buffer-A and B and the pellet, for raw water samples. One hundred microliters of anti-*Cryptosporidium* and 100 µL of anti-*Giardia* Dynabeads were added to the LT tubes and mixed by rotating using at 18 rpm for 1 hour. The LT tubes were placed in a magnetic particle concentrator (MPC) and gently mixed by tilting at 90° angle for 2 minutes. The supernatant was decanted and the LT tubes were removed from the MPC. One millilitre of 1X buffer A was added to the LT tubes to collect the beads. The suspension containing bead-oocysts/cysts complex was transferred into 1.5 mL Eppendorf tubes. These tubes were placed in MPC and rocked for 1 minute. The supernatant was gently removed through aspiration. The magnets were removed, 50 µL of 0.1 N hydrochloric acid (HCl) was added to the tubes and the tube was vortexed for 20 seconds. The tubes were allowed to stand for 10 minutes, the vortexed for 20 seconds and the magnets were inserted. The tubes were allowed to stand undisturbed for few seconds. The beads collected at the back of the tubes and the acidified suspension was transferred into microscope slides containing 10 µL of 1.0 N of Sodium hydroxide (NaOH). This procedure was repeated twice for each tube and transferred into the same microscope slide.

#### Staining procedure (Immunofluorescence assay)

The slides were dried up and fixed using methanol. Twenty five microliters of fluorescein isothiocyanate anti-*Cryptosporidium* and 25 µL of fluorescein isothiocyanate of anti-*Giardia* monoclonal antibodies (FITC-mAb) stains were added onto the slides and incubated at 37 ± 2°C for 45 minutes. The excess FITC-Mab was aspirated. Fifty microliters of 4’6-diamidino-2-phenyindole (DAPI) (50 µL) was added onto the slides and allowed to stand at room temperature for 2 min then the excess was aspirated. Fifty microliters of PBS was added onto the slides and allowed to stand at room temperature for 2 min then the excess was aspirated. The slides were allowed to dry. Diazabicyclooctane (DABCO) mounting medium (8 µL) was used to mount the slides and they were sealed. An Olympus epiflourescence microscope containing the blue filter block was used to examine the stained slides, and to detect and enumerate FITC-mAb labelled *Cryptosporidium* oocysts and *Giardia* cysts at 40x magnification. DAPI stain characteristics were examined at 40x magnification with the UV filter.

## 3. RESULTS

### 3.1 Prevalence and concentrations of *Cryptosporidium oocysts* and *Giardia cysts* in raw influents and final treated effluents

Table 2 provide the results of the prevalence of *Cryptosporidium* oocysts and *Giardia* cysts through the treatment process in WWTP1. A total of 90 samples (36 raw influents, 18 primary treated effluents, 18 secondary treated effluents and 18 final treated effluents) were collected and analysed in WWTP1 over a period of one year. According to these results both *Cryptosporidium* and *Giardia* were detected in 99% (89/90) of the samples collected from WWTP1. As shown in Table 2, *Cryptosporidium* and *Giardia* were detected in 100% (n=36) of raw influent samples. The same result was obtained in primary treated effluent, where both *Cryptosporidium* and *Giardia* was detected in 100% (n=18) of the samples collected. In secondary treated effluent, a total of 18 samples were collected and *Cryptosporidium* was detected in 100% (n=18) of the samples, whilst *Giardia* was detected in 94% (n=18) of the samples. Visa versa results were obtained in the final treated effluent samples, where *Cryptosporidium* was detected in 94% of the samples, whereas *Giardia* was detected in 100% of the samples.

Also shown in table 2 are the results of the prevalence of *Cryptosporidium* oocysts and *Giardia* cysts through the treatment process in WWTP2. A total of 72 samples (18 raw influents, 18 primary treated effluents, 18 secondary treated effluents and 18 final treated effluents) were collected and analysed in WWTP2 over a period of one year. According to these results *Cryptosporidium* was detected in 97% (70/90) of the samples collected, whilst *Giardia* was detected in 99% (71/72) of the samples collected from WWTP1. As illustrated by Table 2, *Crytposporium* and *Giardia* were detected in 100% (n=18) of raw influent samples. The same result was obtained in primary treated effluent, where both *Cryptosporidium* and *Giardia* was detected in 100% (n=18) of the samples collected. In secondary treated effluent, a total of 18 samples were collected and *Cryptosporidium* was detected in 94% (n=18) of the sample, whereas *Giardia* was detected in 100% (n=18) of the samples. In the final treated effluent samples. Both *Cryptosporidium* and *Giardia* where detected 94% (n=18) of the samples collected.

Figure 1 and figure 2 illustrates the mean concentration of both *Cryptosporidium* oocysts and *Giardia* cysts in different treatment stages over 12 months period of the study. According to the results, mean concentration of both *Cryptosporidium* oocysts and *Giardia* cysts were considerable higher in the raw influent treatment stage. Reduction in mean concentration for both *Cryptosporidium* and *Giardia* was observed through the treatment stages with the lowest values obtained in the final treated effluent. WWTP2 had the highest mean concentration for *Crytposporium* (9134 oocysts/ 10 L), whilst for WWTP2 high mean concentration for *Giardia* (13572 cysts/ 10 L).

**Table 2:**
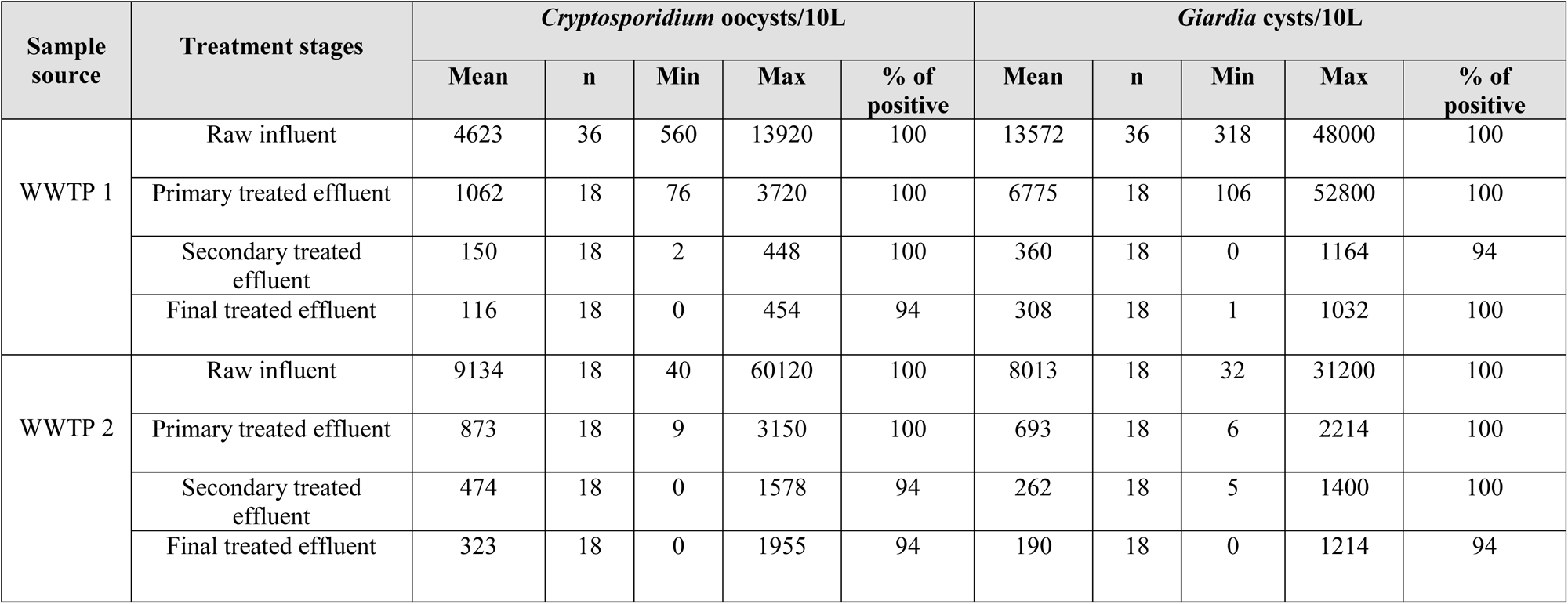
Prevalence; mean, max (maximum), min (minimum) and the percentage number of samples positive for *Cryptosporidium* and *Giardia* in raw influent and the final treated effluent from the two WWTPs.

**Figure 1.**
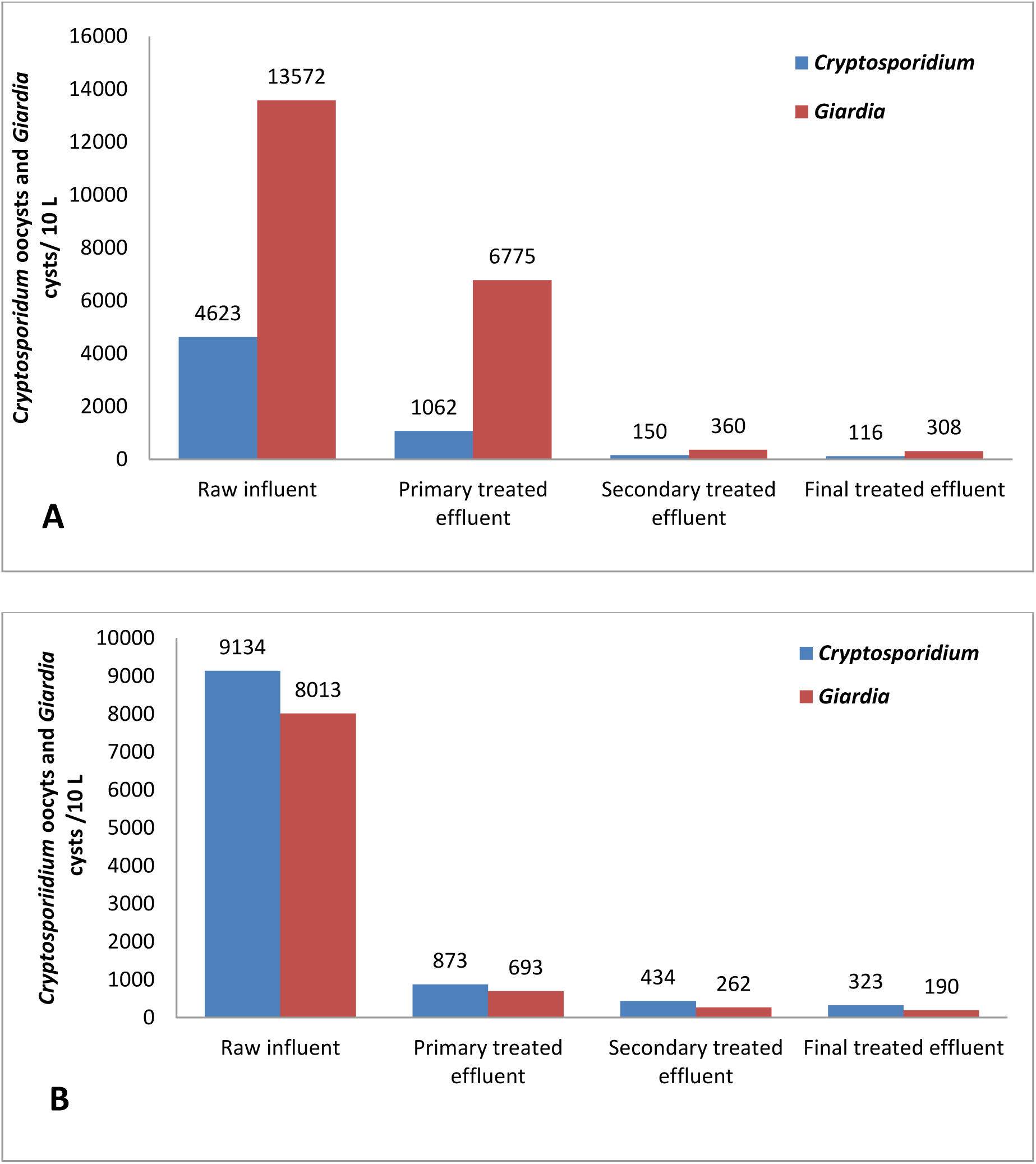
Mean Concentrations of *Cryptosporidium* and *Giardia* in raw influent, primary treated effluent, secondary treated effluent and the final treated effluent at WWPT1 (A) and WWTP2 (B).

**Table 3:** Mean % removal efficiency of *Cryptosporidium* and *Giardia* in WWTP 1 and WTTP 2.

| WWTP | % Removal efficiency |  |
| --- | --- | --- |
|  | *Cryptosporidium* | *Giardia* |
| WWTP 1 | 97.5 | 97.7 |
| WWTP 2 | 96.5 | 97.6 |

## 4. DISCUSSION

This study aimed at determining the prevalence of *Cryptosporidium* and *Giardia* in two South African WWTPs. Using method 1693, *Cryptosporidium* oocysts and *Giardia* cysts were isolated, identified and enumerated in different treatment stages (raw influent, primary treated effluent, secondary treated effluent and final treated effluent) in both WWTPs. These results suggest that *Cryptosporidium* and *Giardia* are one of the predominant pollutants in wastewater treated by the two WWTPs. Moreover, detection of *Cryptosporidium* oocysts and *Giardia* cysts in the final treated effluent indicated that the two WWTPs were inefficient in removing either *Cryptosporidium* oocysts or *Giardia* cysts in the wastewater.

Both *Cryptosporidium* and *Giardia* were detected consistently in all treatment stages (raw influent, primary effluent, secondary effluent and the final treated effluent) at both WWTPs. In raw influent, results obtained revealed that both *Cryptosporidium* oocysts and *Giardia* cysts were consistently detected in all raw influents samples (100% samples) analysed throughout the year. These results indicate a high prevalence of *Cryptosporidium* and *Giardia* in the wastewater treated by the two wastewater treatment plants. The occurrence of *Cryptosporidium* and *Giardia* obtained in this study was higher than that reported by Dungeni and Momba (2010), where *Cryptosporidium* oocysts and *Giardia* cysts were isolated in 50% and 78% of raw influent samples. Robertson and co-workers (2006) also detected *Cryptosporidium* and *Giardia* in 93% and 80% of raw wastewater samples. Also, it is likely that percentages of *Cryptosporidium* and *Giardia* reported in several studies are an underestimation, due to the challenge of low recovery efficiency of the method used. In a study carried out in north-eastern Spain, *Cryptosporidium* was detected in 100% of raw influent wastewater samples (Montemayor et al. 2005). Cheng and co-workers (2009) also obtained both *Cryptosporidium* and *Giardia* in 100% of the samples analysed.

Concentrations of *Cryptosporidium* oocysts and *Giardia* cysts varied throughout the study period with highest concentrations of 60120 oocysts/10L and the lowest concentration of 32 cysts/10L. According to McCuin and Clancy (2006), high variations in concentrations oocysts could be due to several factors such as the 83 levels of infections in the contributing community, the severity of the infection in the community and the size of the community. In the current study industries and abattoirs were also contributors of wastewater. Therefore, levels of *Cryptosporidium* oocysts and *Giardia* cysts in the wastewater discharged by the domestic, industries and abattoirs contributed to the concentration variations. Similar trends where *Cryptosporidium* and *Giardia* concentrations varied between sampling occasions were obtained in a study by Robertson and co-workers (2000). According to Robertson and co-workers (2000), concentrations of *Cryptosporidium* and *Giardia* detected in raw influent are determined by the contributors such as the infected animals and humans in the community served by the wastewater treatment plants (Robertson et al. 2000). The mean concentration of Giardia (13572 cysts/10L) was higher than *Cryptosporidium* concentration in WWTP 1 (*P*=0.01). This phenomenon could indicate that there is a lower prevalence of *Cryptosporidium* in the wastewater contributors (abattoirs, human and industries). On the other hand, results on concentrations of these two protozoans should be interpreted with caution, considering the fact that the method used in this study had low recoveries in the raw influent matrix. Similar results were obtained in a study by Castro-Hermida and co-workers (2010), where mean concentration of *Giardia* was higher than mean concentration of *Cryptosporidium* (Castro-Hermida et al 2010). In WWTP 2, the mean concentration of *Cryptosporidium* (9134 oocysts/10L) was higher than *Giardia* concentration. WWTP 2 treated human associated waste mostly. Therefore, the high mean concentration of *Cryptosporidium* could indicate that *Cryptosporidium* infections were common in the communities served by the WWTP.

Throughout the treatment process, both *Cryptosporidium* and *Giardia* were detected through the period of the study. According to the results, mean concentration of both *Cryptosporidium* oocysts and *Giardia* cysts were considerable higher in the raw influent treatment stage. Reduction in mean concentration for both *Cryptosporidium* and *Giardia* was observed through the treatment stages with the lowest values obtained in the final treated effluent. This trend in concentrations of both protozoans was also observed in study by Cheng and co-workers (2009), where high concentrations of *Cryptosporidium* and *Giardia* were observed in raw influents and low concentrations were obtained in the final treated effluent. Mean removal efficiency ranged from 96.5 to 97.7% for both *Cryptosporidium* oocysts and *Giardia* cysts in both WWTPs. According to Robertson et al (2000), several factors play a role in the removal of parasites by WWTPs. These factors include parasite properties and the properties of the WWTP.

Both WWTPs had high percentage removal efficiency. However, the fact that *Cryptosporidium* and *Giardia* were detected in the final treated effluents discharged by the WWTPs indicated the risk of contamination of the receiving water bodies. This phenomenon is of major concern if the receiving water bodies are being used for domestic purposes by communities living downstream of the WWTPs (Lim et al. 2007). The percentage removal results obtained in the study were similar to those obtained by Dungeni and Momba (2010) and Montemayor et al (2005). Parker et al (1993) obtained a percentage removal efficiency of 80-98% for *Cryptosporidium* oocysts and 98-99% for *Giardia* cysts. Dungeni and Momba (2010) obtained a similar percentage removal efficiency ranging from 67.4% to 98.26% and from 86.81 to 99.96% for *Cryptosporidium* oocysts and *Giardia* cysts respectively.

Results obtained revealed the presence of *Cryptosporidium* oocysts and *Giardia* cysts in final treated wastewater effluent in both WWTPs. In WWTP 1, *Cryptosporidium* oocysts were isolated in most of the treated effluent samples (94%, 0 - 454 *Cryptosporidium* oocysts/10L and 100%, 1 - 1032 *Giardia* cysts/10L). *Cryptosporidium* and *Giardia* were also detected in most treated effluent samples collected from WWTP 2 (94%, 0-1955 *Cryptosporidium* oocysts/10L and 94%, 0-1214 *Giardia* cysts/10L). A significant difference between concentrations of *Cryptosporidium* and *Giardia* were observed in WWTP 1 (*P* = 0.02). However, for WWTP 2, there were no significant difference observed between *Cryptosporidium* and *Giardia*. Moreover, *Cryptosporidium* oocysts and *Giardia* cysts concentrations reported might be an underestimation, since the recovery efficiency data in (not given in this paper) were low and the method was inhibited by unknown matrix components. Nevertheless, detection of *Cryptosporidium* oocysts and *Giardia* cysts in treated effluents from the two WWTPs suggested that the treatment processes used in these wastewater treatment plants were not effective in removing these two protozoans. The prevalence of *Cryptosporidium* and *Giardia* reported in the study were comparable to the results obtained by Montemayor and co-workers (2005), where *Cryptosporidium* oocysts and *Giardia* cysts were detected in 100% of wastewater treated effluent from two WWTPs (Montemayor et al. 2005). The prevalence and concentrations of *Cryptosporidium* and *Giardia* measured in this study were higher than those reported in a study conducted by Dungeni and Momba (2010) in selected South African WWTPs. This could due to differences in the prevalence of *Cryptosporidium* and *Giardia* in the wastewater contributors in different areas and also the differences in the methodology.

## 5. CONCLUSION

The findings of this study revealed that *Cryptosporidium* and *Giardia* were prevalent throughout the treatment process and in the final treated effluent from the two WWTPs. This is an indication that these two WWTPs could pollute the receiving water sources in terms of *Cryptosporidium* and *Giardia*. Pollution of surface water with *Cryptosporidium* and *Giardia* parasites could have serious public health risks when the water is directly used for drinking purposes, irrigation and for recreational purposes. Hence, this study identified a need for *Cryptosporidium* and *Giardia* to be included in the monitoring programmes for treated wastewater effluent. Also, to ensure that the limited South African surface water resources are protected from being polluted by WWTPs, enhancement of the performance of South African WWTPs to remove these protozoan parasites needs serious attention. This process of enhancement will start with the process of determining the main challenges that are hindering the performance of the WWTPs.

